# MicroRNA 21 induces carcinogenesis in hepatic cells by modulating mitochondrial metabolism

**DOI:** 10.1101/2024.04.25.591193

**Authors:** Ashutosh Kumar Maurya, Lincy Edatt, V.B. Sameer Kumar

## Abstract

Mitochondria plays crucial role in cell’s survivability and normal functioning. But in the case of cancer, the mitochondrial machinery (ETC) is altered and glycolytic pathway is activated as an alternate source of energy. The main reason behind the reprogramming of mitochondrial machinery could be mutations in mitochondrial genes or suppression of genes involved in normal functioning of the mitochondria. MicroRNAs could be a key player in modulating the mitochondrial metabolism, as they have targets on various important mitochondrial genes involved in the Electron Transport Chain of the mitochondria. Any alteration in the expression pattern of the mitochondrial genes would directly contribute to the modulation of normal functioning of the mitochondrial machinery. Micro RNA 21 is an oncomiR, located at q arm of the 17^th^ chromosome. MiR 21 has been reported to be involved in many types of cancer. MiR 21 is reported to have targets on many important genes, crucial for cell survivability and proliferation, most of which falls in the category of tumor suppressor genes. With our bioinformatics analysis, we found that miR 21 has targets on important mitochondrial genes involved in the ETC. So, we tried to elucidate the role of miR 21 in modulation of the mitochondrial machinery and role of this alteration in the mitochondrial mechanism in carcinogenesis. Our results revealed that miR 21 have targets on the Cytochrome C Oxidase 1 (Cox1), which is directly involved in the Complex 4 of the electron transport chain. Next we checked the phenotypic effects of this down regulation of Cox1 by measuring the oxygen consumption by the mitochondria and found that O2 consumption goes significantly down in miR 21 over expressing cells. Along with this, we also checked if exosomes from miR 21 overexpressing cancer cells could induce the carcinogenesis in the normal hepatic cells and found that miR 21 accelerates the rate of cellular migration and enhances the colony formation. The results together suggest that miR 21 posses carcinogenic property, possibly by modulating mitochondrial machinery.

## Introduction

Mitochondria are one of the vital organelles in cellular machinery, which play crucial role in cell’s survivability and its normal functioning (Osallame D et.al, 2012). It is also considered as the powerhouse of the cell (Javadov et.al, 2013). The energy required for all the major biochemical processes within the cell at large, is met by the mitochondria (McBride et.al, 2006). Mitochondria play a very significant role in cell growth, division and survivability (Hausenloy et.al, 2010). A number of diseases, including cancer, have been linked to altered mitochondrial machinery (Sheng-Fan Wang et.al, 2023). In the case of cancer, the electron transport chain and Ox-Phos pathways is altered and glycolytic pathway is activated as an alternate source of energy (Hung Wei Lai et.al, 2023).

One of the major reasons behind the reprogramming of mitochondrial machinery could be mutation in mitochondrial genes or suppression of genes involved in mitochondrial metabolism (Naik P. et.al, 2019). In these circumstances, the expression pattern of the key genes involved in the complex 1-4 of ETC is inhibited, reprogramming the mitochondrial system to use a different energy-producing pathway (Garmash E.V, 2022). There are 14 protein-coding genes in the mitochondria and all of which are required in energy production by the mitochondria. The change in expression of any of these genes would significantly contribute to the dysfunction of the mitochondrial function (Alessandra P. et.al, 2013).

MicroRNAs are small non-coding RNAs, which play very important role in regulating various cellular processes (Vaghf A.et.al, 2022). They do so by regulating the expression pattern of the genes, in a fashion similar to RNAi (Jacob O Brien et.al, 2018). MicroRNAs have been reported to regulate the crucial processes at organelle level including mitochondria (Borralho P.M. et.al, 2014). Mitochondrial microRNAs or MitomiRs are miRNAs of nuclear or mitochondrial origin that are translocated to the mitochondria and have a crucial role in regulation of mitochondrial function and metabolism as they have targets on significant genes necessary for the proper operation of the mitochondrial machinery (Purohit et.al, 2020). A normal cell’s transition from normal to malignant may be influenced by the way these mitomiRs are expressed (Hung Yu Lin et.al, 2021).) Since they have targets on genes involved in ATP generation, prerequisite for cellular proliferation, they could alter the expression pattern of these genes and it may lead in hindrance or acceleration in the process of ATP synthesis (Siengdee P. et.al, 2015)

Micro RNA 21 is an oncomiR (Zheng W. et.al, 2018), located at q arm of the 17th chromosome. miR 21 is reported to have targets on many important genes, crucial for cell survivability and proliferation (Huang Y et.al.,2013) most of which falls in the category of tumor suppressor genes (Noraini et.al, 2020). In most of the cancer, miR 21 is highly expressed (Jenike et.al, 2021) and our bioinformatic analysis, showed that miR 21 has targets on important mitochondrial genes involved in the ETC. So, we tried to elucidate the role of miR 21 in modulation of the mitochondrial machinery and role of this alteration in the mitochondrial mechanism in carcinogenesis.

## Methodology

### Cell culture

Hela (cervical cancer cell line), WRL-68 (Human hepatic non cancerous cell line) and HepG2 cells (Hepatic carcinoma cell line) were cultured in DMEM supplemented with 10% FBS, antibiotic-antimycotic solution and L-Glutamine. The cells were maintained under standard culture conditions at 37°C with 5% CO2 and 95% humidity. For experiments, seeding den-sity of 0.4 x 10^4^ cells (96 well), 0.6×10^6^ cells (30mm dish), 0.8×10^6^ cells (60mmdish), 2.2×10^6^ cells (100 mm dish) were used.

### miR 21 cloning

miRNA 21 was cloned in pCMV miR vector between BamH1 and Xho1 restriction sites and successful cloning was confirmed by sequencing.

### Transformation

The competent cells (DH5α) were transformed with miR 21 plasmid by heat shock method where the plasmid was incubated with the competent cells followed by a quick heat shock at 90 degree C for 2 minutes and then immediately transferring it on ice. The transformed cells were then plated on agar plate containing kanamycin.

### Plasmid isolation

A Single colony was picked from the agar plate and grown in the LB broth containing kanamycin. The broth was incubated at 37°C in a shaking incubator. The plasmid was isolated by using Himedia midi kit following manufacturer’s protocol.

### Transfection

HeLa, HepG2 and WRL cells were transfected with plasmids containing micro RNA 21 using PEI reagent. In brief, the cells were seeded with the density of 0.2x105 cells per well, on a 6 well plate to achieve a confluency of 60-70% after 24 hours. 1mg/ml PEI and 300mM sodium chloride was prepared in nuclease free water. About 4 μg of plasmid was added to the 180μl of NaCl and incubated for 5 minutes after a brief vortexing. Following this, 10μl of PEI was added drop by drop to the mixture while vortexing and incubated at room temperature for 15 minutes. Finally this mixture was added drop by drop to each well of the 6 well plate, rocked and incubated at 37°C. After 6 hours of the incubation, the medium was replaced with fresh media and further incubated for 24hours. Following this, the cells transfected with plasmids having fluorescent tags were observed under fluorescent microscope to check the efficiency of transfection.

### Isolation of mitochondria

The mitochondria were isolated from the transfected cells using hypotonic buffer, where the cells were allowed to swell in the buffer for 10 minutes and then break open the cells to release the mitochondria. The cell suspension was then centrifuged at 1300g to remove the cell debris, followed by centrifugation at 12000g to get the mitochondrial pellet. The mitochondrial pellet was suspended in the mitochondrial resuspension buffer.

### Sonication

The mitochondrial pellet was mixed with Lysis buffer and sonicated for 2 minutes at 70% amplitude with 15 sec ON and 30 sec OFF cycle on 4°C. The solution obtained, was centrifuged at 12000g for 10 minutes. The supernatant was collected and protein estimation was done followed by sample preparation for SDS PAGE.

### Protein estimation

Protein level of mitochondria was estimated by Bradford method (Bradford etal,1976). To achieve this, 10μl of sample and 90μl of bradford reagent (50 mg Coomasie Brilliant Blue-G250 in 25ml ethanol and 50ml of phosphoric acid made upto 100ml with water) was added in triplicates in 96 well plate and the absorbance was taken at 595nm by multimode plate reader. The concentration of protein was calculated from the standard plot to BSA with concentration range from 10μg-100μg.

### SDS-PAGE

Protein sample was prepared by mixing of 6x SDS loading dye and boiling it at 90°C for 10 minutes in water bath. The sample was immediately kept on ice and briefly centrifuged before loading on SDS-PAGE gel. The electrophoresis was carried out by using Bio-Rad electrophoresis unit. The protein samples were run through the stacking gel at 80V for 15 minutes and through the resolving gel at 100V at room temperature until the dye reached the end of the gel.

### Western blot analysis

The purity of the mitochondrial pellet was checked by western blot using mitochondria specific antibody (VDAC). Also, the mitochondrial pellet was checked for the nuclear and cytoplasmic contaminants using Histone H3 antibody for Nucleus and Hexokinase HK3 antibody for the cytoplasm.

### RNA Isolation

RNA was isolated from the mitochondrial pellet as well as from the total cell using trizole reagent. Following this, the concentration of the RNA was checked by using nano drop.

### Polyadenylation of RNA

Poly A tail was added to the RNA by Poly A Polymerase enzyme, using manufacturers protocol. This reaction set up was incubated at 37°C for 30 minutes followed by heat inactivation for 5 minutes at 65°C.

### cDNA synthesis

The polyadenylated RNA were used for the synthesis of miRNA 21 specific cDNA by Kang method. Apart from this, total RNA was used to synthesize the cDNA for checking the expression of mitochondrial genes.

### Real Time PCR (qRT PCR)

Quantative real time PCR was performed to check the expression pattern of the microRNA 21 and other mitochondrial genes in mitochondria before and after over expression of miR 21. RNU6B gene was taken as internal refrence gene for microRNA expression study and GAPDH was taken as internal control for mitochondrial genes.

### mRNA stability assay

To elucidate the targeting of mitochondrial genes by miR 21, mRNA stability assay was performed. The cells were transfected with miR 21 using PEI method. 24 hours post transfection the cells were treated with actinomycine D and the cell samples were collected at 0, 1, 3, 6 and 12 hours for gene expression study.

### Oxygraph analysis

To check the phenotypic effects of the down regulation of the mitochondrial genes by miR 21, the oxygraph analysis was performed, where the oxygen consumption level was checked in the miR 21 over expressed samples and compared with the control samples. In brief, the cells were grown in a 6 well plate and transfected with the candidate microRNAs using PEI method. After 48 hours of incubation at 37°C, the cells were trypsinized and the cell pellet was resuspended in respiration buffer. Later, 1ml of the cellular suspension was added to oximeter and oxygen intake reading was recorded for 10 minutes. The readings were used to plot the graph to represent the oxygen consumption by the mitochondria.

### Exosome isolation

The exosomes were isolated from the miR 21 over expressing cells using PEG method. To achieve this the cells were transfected with miR 21 and media was replaced with serum free medium. 48 hours post transfection; the spent media was collected and centrifuged at 2000g for 1/2 hour, to remove cell debris. Following this, the cells were mixed with PEG solution in 1:2 ratios and incubated at 4 degree C overnight. Finally, the solution was centrifuged at 12000g for 1 hour. The exosomal pellet was dissolved in PBS and protein estimation was done. The characterization of the exosomes was done by CD63.

### Cell migration Assay

Cancer cells have the metastatic properties, where they move from its origin to another place and form a secondary tumor. To mimic this in-vitro we perform the cell migration assay with an objective to check, if the exosomes from miR 21 over expressing cancer cells could induce cellular migration of normal Hepatic cells. A scratch was made in the WRL monolayer and treated with exosomes isolated from miR 21 over expressing cancer cells. The cells were allowed to fill the gap formed by the scratch for 48 hours and the images were taken 0, 24, 48 hours respectively and quantified using ImageJ.

### Soft Agar Colony formation assay

To elucidate the role of miR 21 in inducing carcinogenic properties in normal hepatic cells, the colony formation assay was performed where the WRL cells were suspended in the low melting agarose and treated with the exosomes. The cells were incubated at 37°C for 28 days and allowed to form the colonies. The colonies were stained with Coomasie brilliant blue and images were taken from 20 different locations and quantification was done by using ImageJ.

### In-Silico Analysis

miR 21 target prediction analysis and scoring for mitochondrial genes was done using five algorithms i.e. MirBase, miRanda, Target Scan, miRDB and MicroRNA.org. Five highest scored mitochondrial genes (ND6, ATP6, Cyto-B, Cox1, ND4L) were selected for target validation.

### Statistical analysis

All the data in the study were expressed as the mean with the standard error mean of at least three experiments, each done in triplicates. SPSS 11.0 software was used for analysis of statistical significance of difference by Duncan’s One way Analysis of Variance (ANOVA). A value of P<0.05 was considered significant.

## Results

### Levels of miR 21 in cancerous and non-cancerous hepatic cell line

As first step of this study, the cellular and mitochondrial levels of miR 21 in Cancerous and non-cancerous hepatic cell line (i.e. HepG2 and WRL), was checked by qRT-PCR analysis. The result showed that the levels of miR 21 was significantly low in the total cell, as well as in the mitochondria of HepG2 cells, when compared with the cellular and mitochondrial levels in WRL cells. **(Figure 1-Supplementary data)**

**Figure 1:**
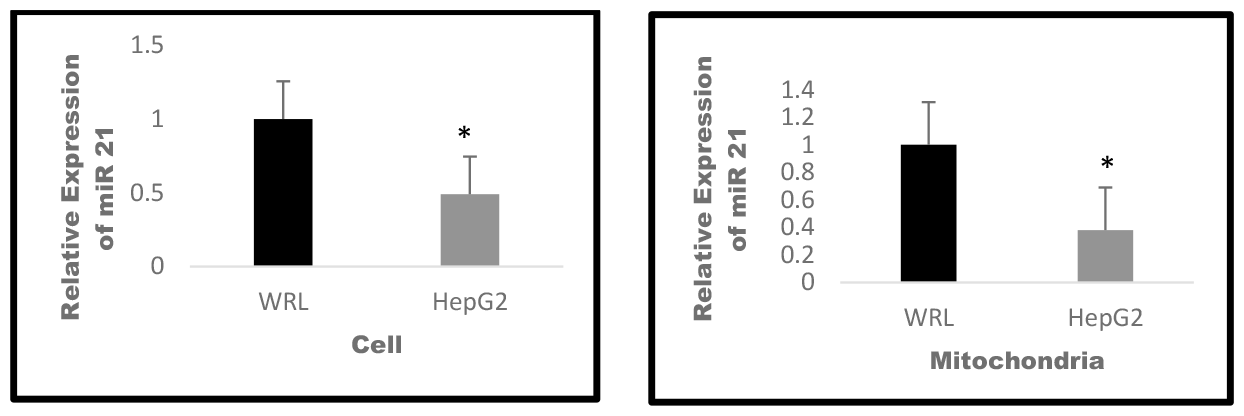
Levels of miR 21 in cytoplasm and mitochondria. RT PCR analysis was performed to check the levels of miR 21 in mitochondrial and cellular fraction of HepG2 cells, keeping WRL as control. The results suggested that level of miR 21 was significantly low in the mitochondria of HepG2 cells. A.) Relative levels of miR 21 in the cells B.) Relative levels of miR 21 in the mitochondria. Results presented are average of three experiments ± SEM each done at least in triplicate, p<0.05. *Statistically significant when compared to control.

### MicroRNA 21 gets enriched in the mitochondria in a preferential manner

As we found that the levels of miR 21 was significantly low in the mitochondria of HepG2 cells, we over expressed miR21 in HepG2 cells to check, if it gets targeted to the mitochondria. The results of the over expression data revealed that, miR 21 gets targeted to the mitochondria in a preferential manner when compared with the mitochondrial and cellular levels in mock transfected cells, with a 4 fold increase in mitochondria as compared to the negligible increase in the cytoplasmic fraction. **(Figure 2-Supplementary data)**

**Figure 2:**
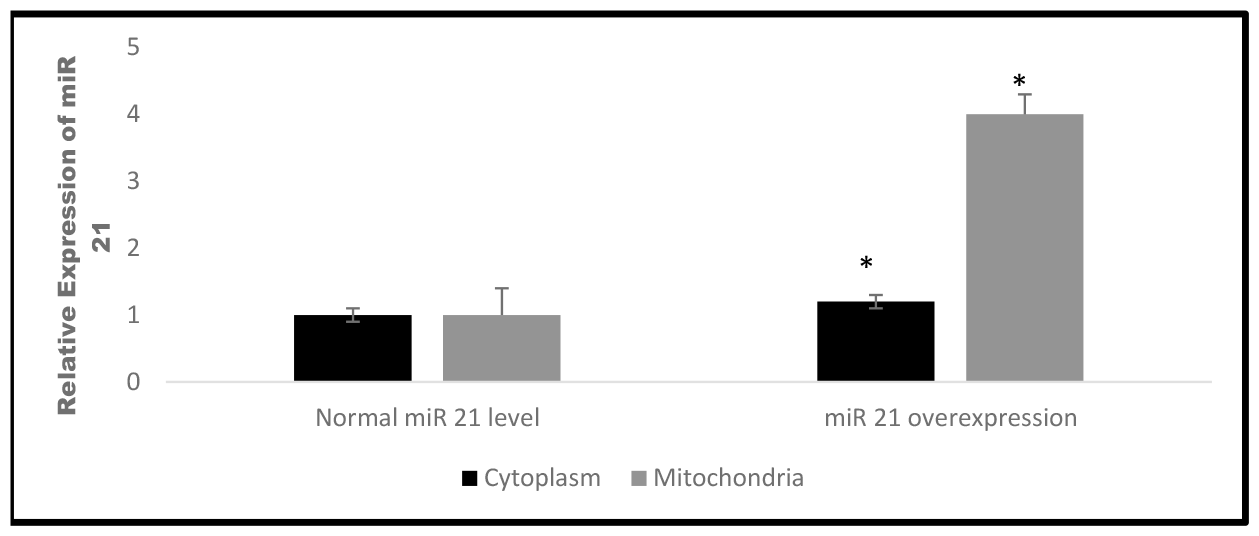
miR 21 gets targeted to the mitochondria in a selective manner. miR 21 was over expressed in HepG2 cells followed by purification of mitochondria & isolation of RNA from mitochondrial as well as cytoplasmic fractions and qRT PCR was then performed to check the levels of miR 21. The results revealed that miR 21 gets targeted to mitochondria in a selective manner. Results presented are average of three experiments ± SEM each done at least in triplicate, p<0.05.*Statistically significant when compared to control.

### miRNA 21 down regulates the expression of its mitochondrial target genes

Upon confirmation of the enrichment of miR 21 in the mitochondria, next we checked the effect of this enrichment on the expression level of mitochondrial target genes. The results revealed that, expression level of all the target genes of miR 21 were significantly lowered under conditions of miR 21 over expression; suggesting that the in-silico target predictions were correct. It was observed that the expression level of Cox1 gene was 3 folds lowered and levels of ATP6, Cyto-B and ND4L were 2.5, 3 and 2.5 folds lowered respectively, when compared with the levels in mock transfected control cells. **(Figure 3-Supplementary data)**

**Figure 3:**
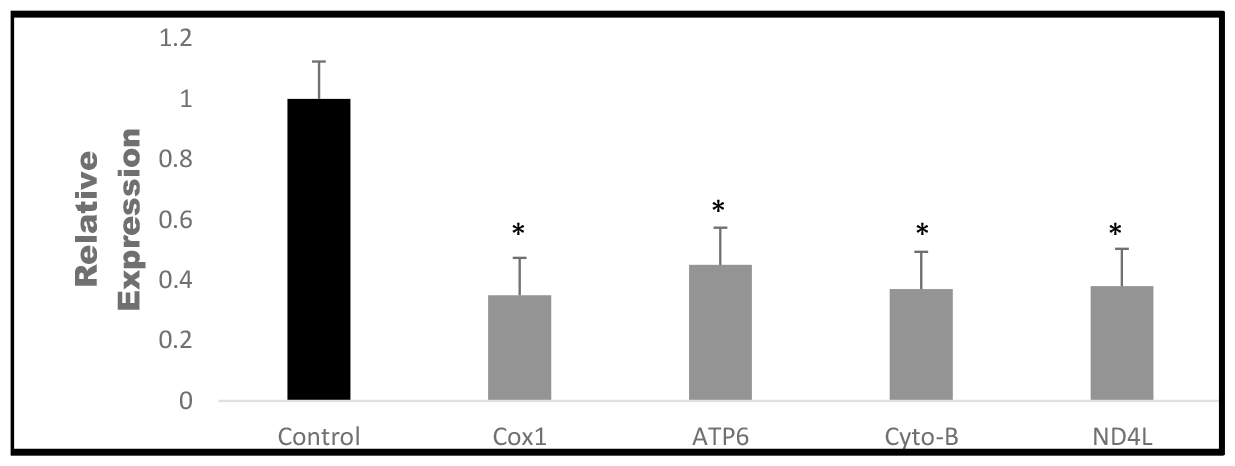
miR 21 down regulates all its target genes. miRNA 21 was over expressed in HepG2 cells and the mitochondria was isolated. RT-PCR analysis was performed to check the impact of increased miR 21 level on the expression pattern of target mitochondrial genes. The result revealed that the expression level of all the mitochondrial target genes of miR 21 went significantly down. Results presented are average of three experiments ± SEM each done at least in triplicate, p<0.05.*Statistically significant when compared to control.

### mRNA stability assay revealed the targeting of mitochondrial Cytochrome C Oxidase (Cox1) by miR 21

With the target validation study, we found that the levels of all the mitochondrial target genes were significantly lowered by miR 21. To further check the targeting of mitochondrial genes by miR 21, we performed mRNA stability assay. The results of the mRNA stability assay revealed that the stability of the mRNA of all the target genes, as evidenced by their levels after treatment with actinomycine, decreased significantly with time under condition of miRNA 21 over expression when compared to the mock transfected control cells. The mRNA stability data suggested that levels of all target genes was lowered by miR 21, most significantly CoX1, suggesting that 3’ UTR of these genes harbor putative miR 21 binding sites. **(Figure 4-Supplementary data)**

**Figure 4:**
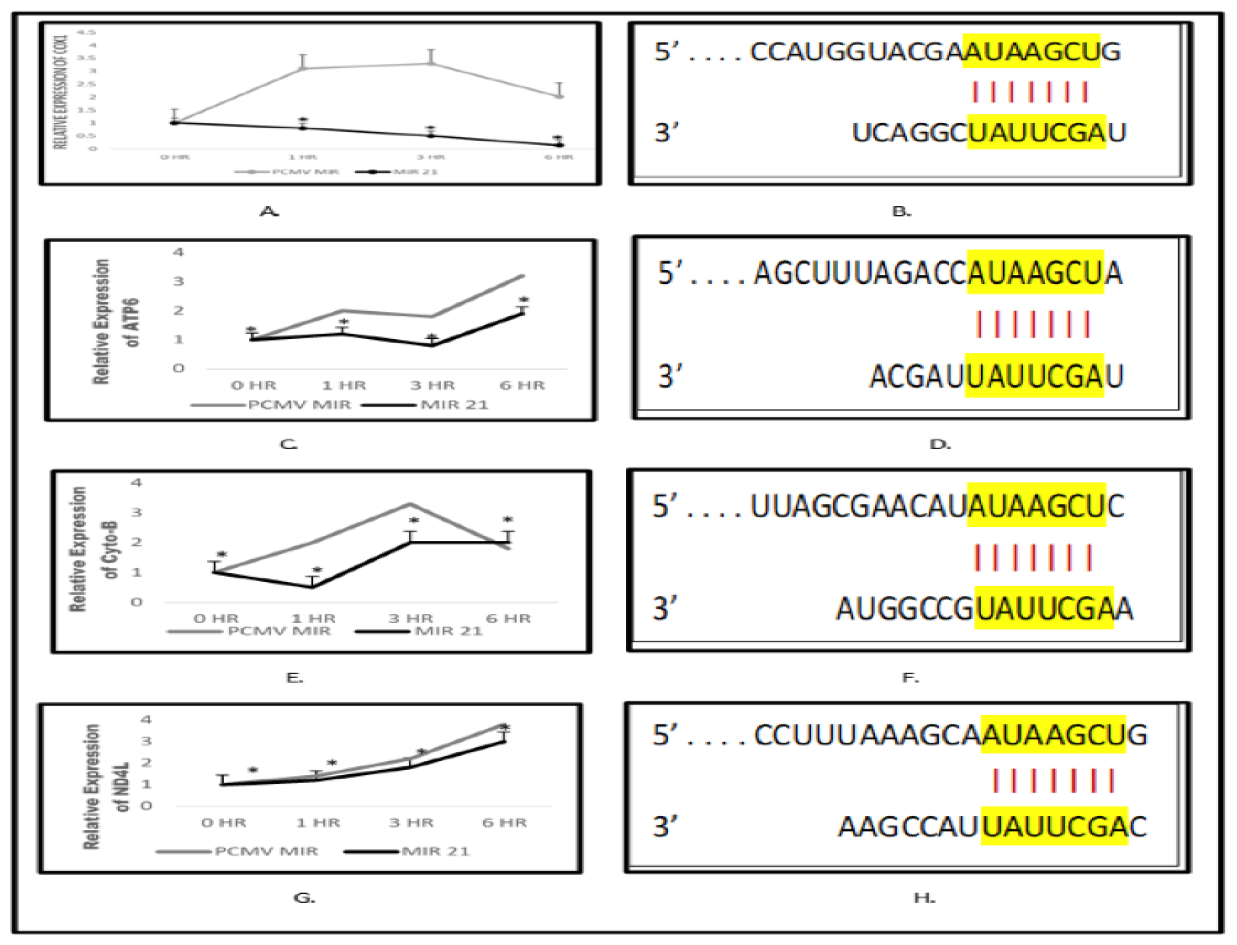
3’ UTR analysis and mRNA stability assay revealed the direct targeting of Cox1 by miR 21. miR 21 was overexpressed in Hepg2 cells and 24 hours post transfection, the cells were treated with actinomycin D and RNA samples were collected at 0, 1, 6 and 12 hours respectively. Following this, RT-PCR analysis was performed to check the expression pattern of target genes. Results revealed that expression of Cox1 went significantly down with time suggesting its direct targeting by miR 21. The expression pattern of ATP6, Cyto-B and ND4L also was altered, but not significant. A.) Relative expression of Cox 1 B.) 3’ UTR analysis of Cox1 C.) Relative expression of ATP6 D.) 3’ UTR analysis of ATP6 E.) Relative expression of Cyto B F.) 3’ UTR analysis of Cyto B G.) Relative expression of ND4L H.) 3’ UTR analysis of ND4L. Results presented are average of three experiments ± SEM each done at least in triplicate, P<0.05.*Statistically significant when compared to control.

### Oxygraph study revealed reduced oxygen consumption by miR 21 over expressing cells

The result of the mRNA stability assay suggested that miR 21 targets the Cox1 gene and since this gene plays very important role in the electron transport chain of the mitochondria, any change in the expression of this gene may alter the normal functioning of the mitochondria. To analyze the phenotypic impact of the down regulation of Cox1 gene on the functioning of the mitochondria, we performed the oxygraph analysis by checking the levels of the oxygen consumption by the HepG2 cells. The result of the oxygraph analysis revealed that the levels of oxygen consumption went significantly down in miR 21 over expressing HepG2 cells when compared with the mock transfected control cells, suggesting its role in altering the mitochondrial machinery by targeting Cox1 gene involved in the complex 4 of electron transport chain of the mitochondria. **(Figure 5-Supplementary data)**

**Figure 5:**
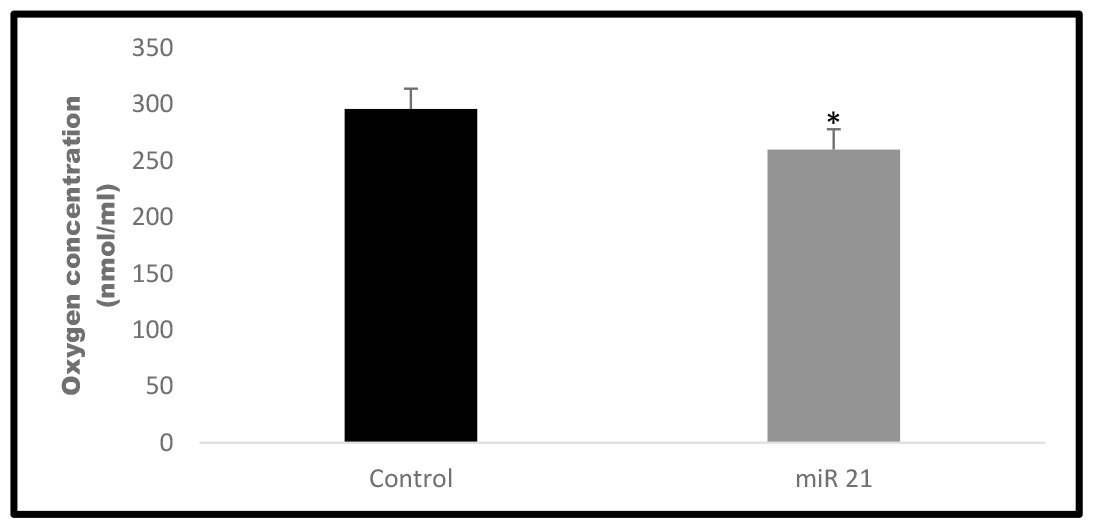
Levels of oxygen consumption by the cells went significantly down, when miR 21 was over expressed. miR 21 was over expressed in HepG2 cell line and 24 hour post transfection, cells were harvested and oxygraph analysis was performed. The results revealed a decrease in O_2_ consumption by the mitochondria of miR 21 overexpressing cells when compared with the mock transfected cell. Results presented are average of three experiments ± SEM each done at least in triplicate, p<0.05.*Statistically significant when compared to control.

### MicroRNA 21 enhanced cellular migration in hepatic cells

With our target validation and mRNA stability assay results, we found that miR 21 get targeted to the mitochondria and down regulate the expression of Cox1 gene, important in ETC and it resulted in the reduced oxygen consumption by the HepG2 cells, probably due to the altered mitochondrial metabolism. So, we next checked the impact of this altered mitochondrial machinery in carcinogenesis. To achieve this, we performed the scratch assay using non-cancerous & cancerous hepatic cells (WRL & HepG2) and treated it with the exosomes isolated from miR 21 over expressing cancer cells and mock transfected cells. The results shown an increase in the rate of cellular migration of non-cancerous hepatic cells (WRL), when treated with the exosomes isolated from miR 21 over expressing cancerous cell line (HeLa). **(Figure 6 & 7 -Supplementary data)**

**Figure 6:**
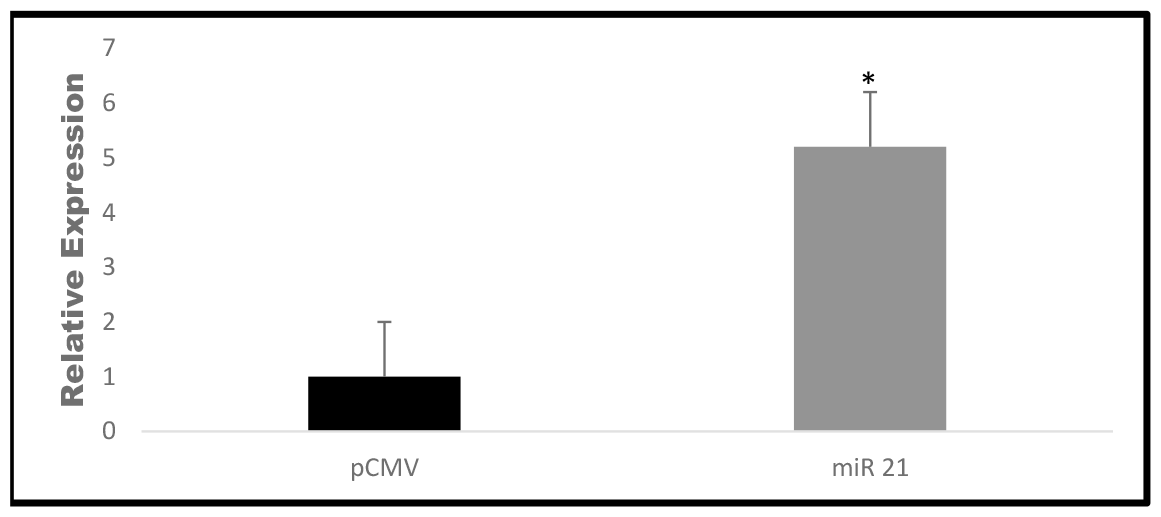
Exosomes isolated from miR 21 over expressing cells were enriched with miR 21. Real time PCR analysis was performed to check the levels of miR 21 in the exosomes isolated from miR 21 over expressing cells and compared with the exosomes isolated from the mock transfected control cells. The results revealed that exosomes isolated from microRNA 4263 over expressing cells, shown 5 fold higher level of microRNA 21. Results presented are average of three experiments ± SEM each done at least in triplicate, p<0.05.*Statistically significant when compared to control.

**Figure 7:**
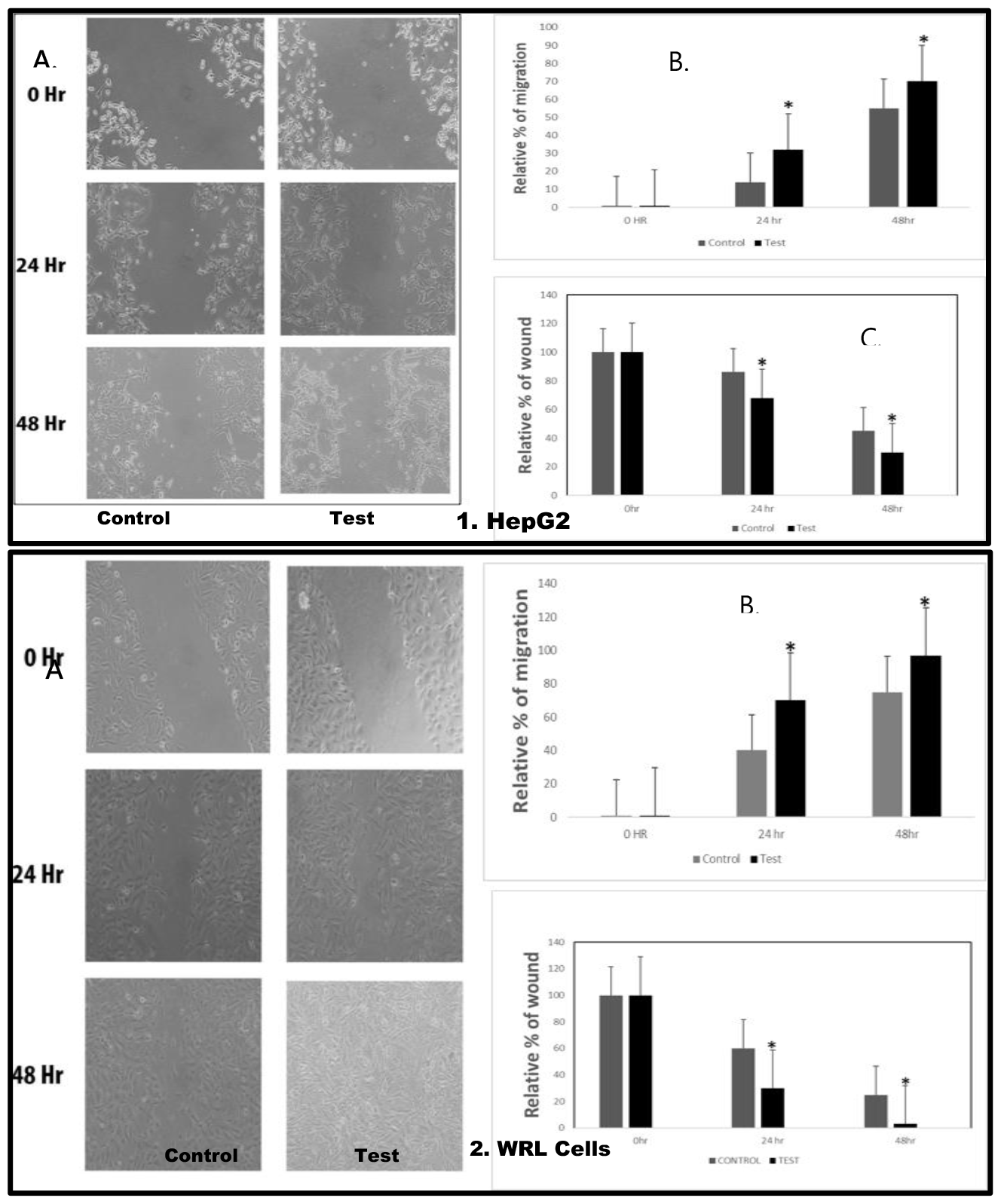
miR 21 enhances the cellular migration when treated with exosomes isolated from microRNA 21 over expressing cells. Cells were grown in a monolayer and a scratch was made, followed by exosome treatment. The microphotographs were taken at 0, 24, and 48 hours. The distance/gap covered by the cells with time was estimated by image-J software. The result revealed that the rate of the cellular migration got escalated when treated with exosomes enriched with miR 21. 1A.) Microphotograph of cell migration pattern (HepG2) with respect to time 1B.) Relative percentage of migration at 0, 24 & 48 hours 1C.) Relative percentage of wound healing at 0, 24 & 48 hours. 2A.) Microphotograph of cell migration pattern (WRL) with respect to time 2B.) Relative percentage of migration at 0, 24 & 48 hours 2C.) Relative percentage of wound healing at 0, 24 & 48 hours. Results presented are average of three experiments ± SEM each done at least in triplicate, p<0.05.*Statistically significant when compared to control.

**Figure 8:**
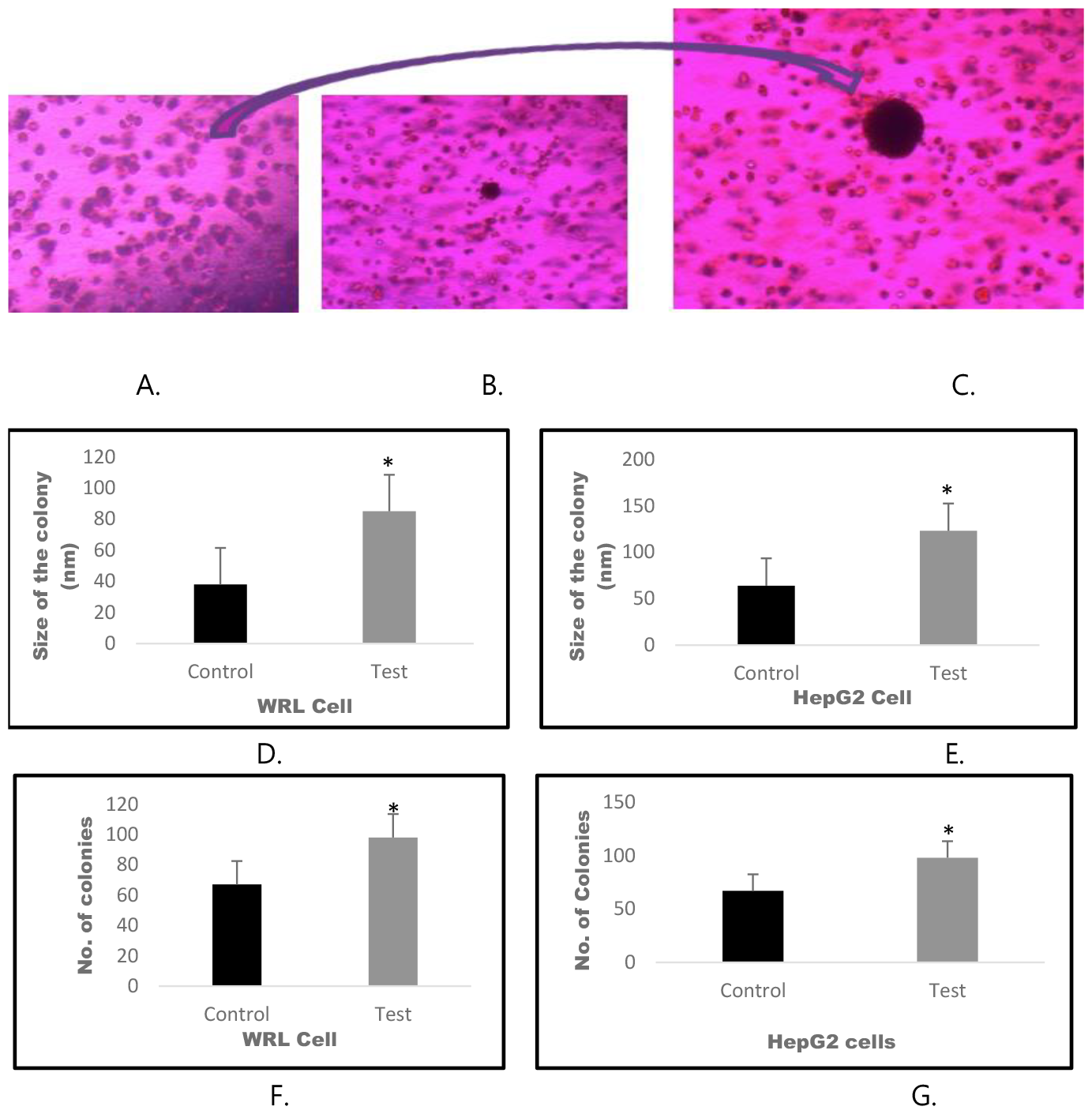
miR 21 enhances the tumor size when treated with exosomes isolated from microRNA 21 over expressing cells. HepG2 and WRL colonies were treated with exosomes isolated from the miR 21 overexpressing cells and allowed to grow for 28 days. Following this, the microphotographs were taken at 20 different regions and 20 colonies from each region was taken for the size estimation by image-J software. The number of colonies were counted manually from 20 regions to estimate the number of tumor colonies. The results revealed that the size as well as the number of colonies got enhanced in HepG2 and WRL cells, when treated with exosomes enriched with miR 21. A.) Representative microphotograph of WRL cell colony B.) Representative micro photograph of HepG2 colony C.) Representative micro photograph of single colony. D.) Comparative colony size of WRL cells. E.) Comparative colony size of HepG2 cells F.) Number of colonies of WRL cells, when compared with control G.) Number of colonies of HepG2 cells, when compared with control. Results presented are average of three experiments ± SEM each done at leastin triplicate, p< 0.05.*Statistically significant when compared to control.

### MicroRNA 21 induces colony formation in hepatic cells

In addition to cellular migration assay, we also performed the colony formation assay to check the carcinogenic ability of miR 21 in normal hepatic cells. Here, we treated the WRL cell colonies with the exosomes isolated from miR 21 over expressing cells or mock transfected cells. After 28 days, the colonies were micro photographed for 20 different regions and 20 different colonies were selected from each region to measure the size of the colonies by image-J software. Colony formation assay result, showed an increase in the size of the colonies when treated with the exosomes enriched with miR21, when compared with colonies treated with exosomes from the mock transfected cells. **(Figure 4-Supplementary data)**

## Discussion

Mitochondria, the powerhouse of the cells, are the primary organelles that supply cellular energy (Zhang et.al, 2021). In addition to producing ATP, mitochondria are crucial for the complete metabolic process of cells, which includes iron metabolism, fatty acid oxidation, amino acid metabolism, formation of reactive oxygen species (ROS), and calcium homeostasis (Nakhle J et.al.,2020, Duarte et. al, 2014, Palmeira et. al, 2007). Mitochondrial dysfunctions are associated with various human diseases, such as diabetes, cardiac diseases, and cancer (Monsalve M et. al, 2007, Sripada L et. al, 2012).

Mitochondrial malfunction results due to inadequate number of mitochondria, or reprogrammed electron transport and adenosine triphosphate synthesis machinery (Duarairaj S et. al, 2020). Occurrences of mitochondrial dysfunction could be due to the lower expression of the mitochondrial genes, crucial for its survival and function. MiRNAs are a class of small, endogenous, noncoding RNAs of 22nt length, which regulates the gene expression by binding to 3 untranslated region of target mRNA (Akhtar M et. al, 2016).

The microRNAs, possessing ability to change the expression pattern of the mitochondrial genes and modulate the mitochondrial system are called MitomiRs (mitochondrial microRNAs). mitomiRs can be transcribed either from the nuclear or mitochondrial genome and localize in mitochondria (Mora Morri et. al, 2018). Mitochondrial miRNAs are known to serve as primary regulators of mitochondrial function (Song R. et. al, 2019) by controlling mitochondrial activity, either by targeting cytoplasmatic mRNAs or by acting inside the mitochondria (David R et. al, 2023). Mitochondrial microRNAs play very important role in carcinogenesis as they have targets on the crucial genes involved in the ETC. Various studies have shown that the mitochondrial machinery is altered in the advance stages of the cancer and glycolytic pathway is activated as an alternate source of the energy production. This modulation of the mitochondrial machinery may result either by mutation or suppression of the genes involved in the ETC and Ox-Phos pathways (Palmeria M et. al, 2007).

MicroRNA-21, located at chromosome 17q23.2, 21 is an abundantly expressed miRNA in mammalian cells, whose upregulation is associated with numerous type of cancers (Feng Y et. al, 2011, Fulci V et. al, 2007). It is one of the earliest identified cancer-promoting ‘oncomiRs’, targeting various tumor suppressor genes linked with proliferation, migration and invasion (Feng Y et. al, 2016). Studies suggests that the knockdown of miR-21 in hepatic cells, increased the expression level of tumor suppressor gene PTEN, which is a direct target of miR-21, and decreased the tumor cell proliferation and migration (Meng F et. al, 2007). MicroRNA-21 has been found to be the most commonly upregulated miRNA in solid tumor including HCC (Volinia S et. al, 2006).

The primary goal of our study was to elucidate the role of miR21 in modulating the mitochondrial machinery and effects of this reprogramming on carcinogenesis. Since, miR 21 is a nuclear coded microRNA, we first checked its levels in the mitochondria of HepG2 cells keeping WRL as control, by qRT PCR analysis and found that levels of miR 21 was very low in the mitochondria of HepG2 cells. Several recent studies suggests that nuclear coded micro RNAs gets targeted to the mitochondria (Huaping L et. al, 2016). We over expressed miR 21 in cells and qRT PCR data revealed that, miR 21 got enriched in the mitochondria 4 folds higher than the cytoplasm, in a selective manner.

Bioinformatic analysis, revealed that miR 21 has targets on 5 important mitochondrial genes i.e. ND4L, ND6, Cyto B, Cox1, ATP6, crucial for complex 1, 3, 4 and 5 of the electron transport chain respectively. To check it further, we performed a target validation study and analyzed, whether miR 21 could downregulate its target genes in mitochondria. The qRT PCR results suggested that miR 21 down regulated all of its five target genes, significantly, Cox1 gene. Next, we performed mRNA stability assay, to further analyze the targeting of Cox1 gene by miR21, and the result revealed that Cox1 gene gets targeted by miR 21 and falls in line with the our 3’ UTR analysis which revealed that 3’ UTR of Cox1 gene harbor putative miR21 binding sites.

Since, Cox1 gene plays important role in the electron transport chain, its down regulation would definitely alter the mitochondrial functioning. So, next we evaluated the phenotypic effect of this down regulation by oxygraph analysis. Here we measured the oxygen consumption by the mitochondria of miR21 over expressing cells and compared it with the normal cells. The results suggested that the oxygen consumption by miR 21 over expressing cells went significantly down, when compared with the control cells. This suggested that miR21 alters the mitochondrial machinery, possibly by targeting Cox1 gene involved in complex 4 of the electron transport chain. Dysregulation of mitochondria is involved in various liver disease, especially Hepatocellular carcinoma (HCC) (Andreas k et. al, 2011), which is one of the most common malignancies, ranking second worldwide with regard to tumor related death (Jemal A et. al, 2011). Aberrant expression of miR-21 could contribute to HCC growth and spread by modulating PTEN expression and PTEN-dependent pathways involved in mediating phenotypic characteristics of cancer cells such as cell growth, migration, and invasion (Meng F et. al, 2007). Studies on clinical data showed that miR-21 was significantly upregulated in both HCC tissues and serum (Karakatsanis A et. al, 2013, Yoon J et. al, 2018,), when compared with normal adjacent liver tissues (Wang W et. al, 2014).

Under the light of the fact that, miR 21 was found less in normal liver tissues and it can increase tumor cells migration (Liu C et. al, 2010, Asangani I et. al, 2008), as well as could promote cellular proliferation (Hao X et. al, 2018), we next tried to check if miR 21 can induce migratory and colonogenic properties in normal hepatic cell lines. To achieve this, we isolated exosomes from miR 21 over expressing cancer cells and performed cell migration and colony formation assay in hepatic cell line. The results of colony formation assay revealed an increase in number and size of the colonies in cells treated with the exosomes enriched with miR21, when compared with the cells treated with exosomes from mock transfected cells. Further, cell migration assay also revealed an increase in the rate of cellular migration when treated with exosomes enriched with miR 21 when compared to control.

Our study thus revealed that miR 21 gets targeted to the mitochondria in a selective manner and down regulated the Cox1 gene involved in the complex 4 of the Electron Transport Chain. This down regulation resulted in decreased O2 consumption by the mitochondria, suggesting a possible modulation in the mitochondrial machinery. The cell migration and the colony formation assay results revealed that miR 21 could induce the carcinogenic properties in the normal hepatic cells. All together, our results suggest that the oncogenic property of miR21 in hepatocellular carcinoma possibly involve the alteration of mitochondrial machinery.

## Acknowledgment

We acknowledge Indian Council of Medical Research, Ministry of Health, Govt. of India, for the financial assistance in form SRF and Kerala State Council for Science Technology & Environment, Govt. of Kerala, for fellowship in the form of JRF and SRF to Mr. Ashutosh K. Maurya. We also acknowledge Central University of Kerala for providing all the necessary facilities to carry out this research work.

## Author Contributions

The authors confirm contribution to the paper as follows: Study conception and design: VBSK, Bioinformatics and wet lab work: AKM. Cloning of miR 21: LE. mRNA stability assay (sample preparation): RP. All authors reviewed the results and approved the final version of the manuscript.

## Supplementary data

